# Coordinated neural, metabolic and muscular transcriptomic signatures associated with mite-biting behavior in honeybees (*Apis mellifera* L.)

**DOI:** 10.64898/2026.05.27.728268

**Authors:** Rameshwor Pudasaini, M. Catherine Farrell, Hongmei Li-Byarlay

## Abstract

Biting behavior is an important natural defense mechanism in honeybees (*Apis mellifera*) against *Varroa destructor*. Significant variation in this behavior exists across genetic lines of honeybees, with certain colonies exhibiting higher mite-biting activity than others. Selective breeding for enhanced biting behavior provides a promising strategy for sustainable mite control and colony resilience. However, successful implementation of such breeding programs requires a comprehensive understanding of the genomic mechanism underlying this trait. In this study, RNA-seq analysis of mandible transcriptomes of 1-day and 8-day old worker honeybees from high mite biting (HB) and low mite biting (LB) colonies were performed. A total of 9,345 genes (97.30%) showed a significant differential expression between LB and HB honeybees across different ages (one-way ANOVA, FDR < 0.05). Comparison of LB vs. HB workers collected on day 1 detected 166 down-regulated and 403 up-regulated genes, whereas workers collected on day 8 identified 82 down-regulated and 46 up-regulated genes. Furthermore, the Weighted Gene Coexpression Network Analysis (WGCNA) exhibited the brown and yellow modules with significantly higher expression in HB compared with LB on both day 1 (FC = 1.52, FDR = 0.0019 and FC = 1.47, FDR = 2.7 × 10^-4^, respectively) and day 8 (FC = 1.27, FDR = 0.056 and FC = 1.16, FDR = 0.073, respectively). Gene Ontology (GO) enrichment analysis identified over-representation of biological processes involved in muscle contraction, chitin binding, neural signaling, oxidative phosphorylation, neuron development, electron transport chain, mitochondrial ATP synthesis, stress response, sensory perception and metabolic processes. Kyoto Encyclopedia of Genes and Genomes (KEGG) pathways analysis identified significant enrichment of various pathways including oxidative phosphorylation, cytoskeleton-related pathways, carbon metabolism, motor proteins, ribosome-associated pathways, and citrate cycle (TCA cycle). The present findings demonstrate mite-biting behavior is associated with coordinated activation of neural, energetic and muscular system rather than a single molecular mechanism. These findings provide a basis to improve honeybee health, enhance resistance to *Varroa* and other ectoparasite, and eventually support sustainable beekeeping and agricultural pollination systems.

## Introduction

Honeybees (*Apis mellifera* L.) play a critical role in global agriculture through pollination, contributing to both ecosystem stability and crop production (Khalifa et al., 2021; Siviter et al., 2023). It is the most important managed pollinator for global and domestic food security (Khalifa et al., 2021; Walsh & Simone-Finstrom, 2024). Honeybees pollinate about US$ 15 billion worth of crops in the United States each year, including more than 130 types of fruits, nuts, and vegetables (USDA, 2026). However, due to multiple interacting factors including both biotic and abiotic stress, such as parasites, pesticides, habitat loss and disease, honeybee populations are declining globally (Willcox et al., 2023).

*Varroa destructor* is considered the most serious threat to honeybees in North America and worldwide (Morfin, Goodwin, et al., 2023; Traynor et al., 2020). The *Varroa* mite feeds on the fat tissue of both immature and adult honeybees that resulted in depriving energy reserve in honeybees and reducing their ability to mount immune responses (Ramsey et al., 2019). Apart from that they also they are also vector of several honeybee viruses including deformed wing virus complex (DWV), Israeli Acute Paralysis Virus (IAPV), Acute Bee Paralysis Virus (ABPV), and Kashmir Bee Virus (KBV) (Lamas et al., 2026; Traynor et al., 2020). These viruses interact with other stressors and often result in synergistic effects on colony health (Hsieh & Dolezal, 2024). These virus infected honeybees also display different morphological deformities such as deformed wings, shortened abdomens, and reduced body size (Lamas et al., 2026; Ullah et al., 2021). Similarly, virus infected honeybee colonies may also have higher chance of colony collapse due to reducing workers’ lifespan, brood mortality and impair in their social behavior (Geffre et al., 2020; Ullah et al., 2021). Therefore, beekeepers need to heavily depend on synthetic acaricides to control *Varroa* infestations in honeybee colonies (Lester, 2023; Qadir et al., 2021). However, in *A. mellifera* colonies, *Varroa* are reported developing resistance to acaricides globally (Bertola & Mutinelli, 2025; Roth et al., 2021).

To combat with *Varroa,* honeybees also exhibit complex and coordinated defensive behavior, such as hygienic and *Varroa*-sensitive hygiene (VSH) behavior, grooming behavior including biting behavior and brood recapping behavior (Morin & Giovenazzo, 2023). Among these, biting behavior has emerged as an important natural defense against *Varroa* (Ulgezen et al., 2021).Honeybee workers use their mandibles to bite, damage and remove mites from themselves or other colony members (Li-Byarlay et al., 2025; Smith et al., 2021). The biting behavior vary among genetic lines, with some colonies exhibiting high biting activities while others display low levels (Akongte et al., 2023; Li-Byarlay et al., 2025). A study reported that *V. destructor* populations grew significantly less in high mite biting colonies compared to less mite biting colonies (Morfin et al., 2020). Studies also show that this mite biting trait has a heritable genetic basis and can be enhanced through selective breeding (Hunt et al., 2016; Li-Byarlay et al., 2025; Morfin et al., 2020; Smith et al., 2021). Selection and breeding of honeybee colonies with high biting behavior to *Varroa* could help to dependence on synthetic acaricides. Therefore, for successful selection and breeding, it is mandatory to understand both behavior assays and genomic underlaying mechanism associated with high biting behavior.

Age-related behavioral transitions in worker honeybees influence overall colony health and resilience (Smith et al., 2022; Vance et al., 2009). These changes are strongly influenced by genetics, such as the shift from nursing to defense or foraging (Whitfield et al., 2006). Honeybees performing different age-associated tasks exhibit distinct molecular and physiological profiles, including differences in biting behavior based on different ages (Whitfield et al., 2003). Transcriptomic analysis provides a powerful approach to profile gene expression genome-wide and to uncover the genetic and molecular pathways associated with complex behaviors (Bresnahan et al., 2023; Morfin, Harpur, et al., 2023). In honeybees, transcriptomic studies have successfully linked behavioral phenotypes to changes in neural, immune and metabolic pathways, highlighting the utility of this approach for understanding the biological basis of social behavior (Bresnahan et al., 2023; Chandrasekaran et al., 2011). However, comprehensive analysis of comparing low-and high biting bees across different age groups is still lacking.

In this study, we performed a comparative transcriptomic analysis of low mite biting (LB) and high mite biting (HB) honeybees collected on day 1 and day 8 to identify differentially expressed genes and pathways associated with biting behavior. By integrating behavioral phenotypes with gene expression profiles, our study aims to reveal the molecular drivers of this defensive behavior, providing insights that could inform selective breeding strategies for improved colony health and resilience.

## Materials and Methods

### Honeybee collection

Based on mite biting behavior, previously classified as LB and HB colonies at the Central State University, Wilberforce, Ohio Apiary were used in this study. Three individual day-one workers were collected from each of the LB and HB colonies (*n* = 6) and kept each of them in a 1.5 mL centrifuge tube (ThermoFisher Scientific). Each tube including collected a worker was freshly frozen in liquid nitrogen and stored at 80 °C freezer prior to the RNA extraction. From the same LB and HB colonies, some one-day-old workers of each LB and HB colonies were individually painted with a blue Sharpie® paint marker on their thorax, then released back to their hive. On the 8^th^ day of marking, 3 marked individual workers were collected from each of the LB and HB colonies (*n* = 6) and preserved as described above. Overall, this study consisted of 2 treatment groups (LB and HB colonies) with 2 time points, with 3 worker honeybees collected from each colony at day 1 and day 8 (*n* = 12).

### RNA extraction, library preparation and Illumina sequencing

Total RNA was extracted using whole honeybee samples using the RNeasy Mini Kits (Qiagen, Hilden, Germany) with following the manufacturer’s protocol. RNA concentration and purity were measured using a NanoDrop spectrophotometer (Thermo Fisher Scientific, USA) and RNA integrity was determined using an Agilent 4200 TapeStation system (Agilent Technologies, USA). Next, Qiagen’s QIAseq Stranded mRNA Enrichment Kit (Qiagen, Hilden, Germany) in combination with the QIAseq Low Input RNA Library Kit (Qiagen, Hilden, Germany) were used for mRNA libraries preparation. Both N6-T RT and ODT-RT primers for reverse transcription were used during mRNA libraries preparation. Qubit fluorometer (Thermo Fisher Scientific, USA) was used for quantifying RNA and a total of 200 ng of input RNA per sample was used for library construction. RNA fragmentation was performed for 3 minutes, followed by first-and second-strand synthesis and adapter ligation according to the manufacturer’s protocol. Unique dual indexes were added using the QIAseq UX96 Index Kit UDI-B (Qiagen, Hilden, Germany). A total of 18 PCR cycles were used for libraries amplification, and further purified, determined quality checked prior to sequencing with Agilent 4200 TapeStation system (Agilent Technologies, USA) and Qubit Fluorometer (Thermo Fisher Scientific, USA). Final libraries were pooled and sequenced on an Illumina NovaSeq X Plus platform (Illumina Ibc., San Diego, CA, USA) using paired-end 150 bp reads to generate high-depth mRNA expression data.

### Quality check, alignment and count generation

#### Reference sequences

The *A. mellifera* transcriptome file “GCF_003254395.2_Amel_HAv3.1_rna.fna.gz” from NCBI was used for quasi-mapping and count generation (https://www.ncbi.nlm.nih.gov/datasets/genome/GCF_003254395.2/). The gene model file “GCF_003254395.2_Amel_HAv3.1_genomic.gff.gz” was used for traditional alignment and gene counting as well as to generate a transcript-gene mapping table for obtaining gene-level counts.

### Quality check on raw data and alignment

Raw individual reads underwent quality control using FASTQC (version 0.11.9) and low-quality based and adapter sequencing were trimmed before further analysis (Andrews, 2010). The reaming reads were then results summarized into a single HTML report by using MultiQC (version 1.12) (Ewels et al., 2016).

Reads were aligned using STAR (version 2.7.11b) (Dobin et al., 2013), as it yielded better alignment rates than Salmon (version 1.10.1) (Patro et al., 2017). Percentage reads that overlapped any exon of a gene (in.a.gene) ranged from 83.9% to 89.8%.

### Sample quality control

#### Normalization

The trimmed mean of M values (TMM) normalization (Robinson & Oshlack, 2010) was performed with edgeR package (version 4.6.2) (Robinson et al., 2010) with using the assumption of *most genes do not change* to calculate a normalization factor for each sample to adjust for such biases in RNA composition. In this dataset, TMM normalization factors fluctuate between 0.61 and 1.23.

### Filtering, clustering and remove unwanted variation (RUV) analysis

The detection threshold was set at 1 count per million (CPM) in at least 3 samples, which resulted in out of a total 12,388 genes (Amel_HAv3.1 (+ endemic bee viruses)), 2,784 genes being filtered out, and leaving 9,604 genes to be analyzed for differential expressions that contain 99.96% of the reads. After filtering, TMM normalization was performed again, and normalized log2-based CPM (logCPM) were calculated using CPM () function with edgeR (version 4.6.2) (McCarthy et al., 2012) function with *prior.count = 2* to help stabilize fold-changes of extremely low expression genes.

Multidimensional scaling in the limma package (version 3.64.1) (Ritchie et al., 2015) was used as a sample quality control step to check for outliers or batch effects. The normalized and filtered logCPM values of the top 5,000 variable genes were chosen to construct the multidimensional scaling plot.

The RUVSeq was used to correct a batch effect in dimension 2 (23.6% of total variability) of multidimensional scaling (MDS) clustering (Risso et al., 2014). The RUV factor started with 1 to determine if it would remove the unwanted variation seen in dimension 2 of MDS and it did accomplish this. The limma-voom method (Law et al., 2014) was used to calculate a one-way ANOVA test across all treatment groups and counted the number of genes with a raw p-value < 0.05. The clustering of treatment group replicates has improved greatly after RUV analysis, Dimension 1 (62.8%) separates “day 1” samples from the “day 8” samples, and Dimension 2 (17.4%) separates LB from HB samples. The RUV factor is added to the statistical models used in the sections following this when calculating the difference between groups.

### Differential gene expression analysis

Differential gene expressions (DGEs) analysis was performed using the limma-trend method (Chen et al., 2016) with a one-way ANOVA test across all treatment groups. Multiple testing correction was done using the False Discovery Rate method (Benjamini & Hochberg, 1995) on the one-way ANOVA raw p-values.

### Pairwise Comparisons

To perform statistical tests between pairwise combinations of groups, LB vs. HB across day 1 vs. day 8 comparisons were conducted. Additionally, statistical tests of the interaction between them were also performed. The names of the comparisons and their mathematical formulations are presented below. The first group listed is in the numerator of the fold-change and the second group listed is in the denominator.

Effect of day on gene expression in LB (Day 8 vs. Day 1) = LB_8 - LB_1

Effect of day on gene expression in HB (Day 8 vs. Day 1) = HB_8 - HB_1

Effect of treatment with day 1 (HB vs. LB) = HB_1 - LB_1

Effect of treatment with day 8 (HB vs. LB) = HB_8 - LB_8

Interaction between day and treatment = (HB_8 - HB_1) - (LB_8 - LB_1)

### Weighted Gene Coexpression Network Analysis (WGCNA) analysis

A WGCNA analysis was employed to determine the broad-scale patterns of expression across all samples (Zhang & Horvath, 2005).The normalized log_2_ expression values were used that had the 1 surrogate variable removed and selected 9,345 genes with a one-way ANOVA (False discovery rate (FDR) *p*-value < 0.25). The estimated power for soft thresholding was 6 and the blockwiseModules function was used with default variables except maxBlockSize = 20000, networkType = "signed hybrid", corType = "bicor", maxPOutliers = 0.05, minModuleSize = 20, mergeCutHeight = 0.4.

### Functional enrichment analysis

Further functional enrichment analysis was performed on each WGCNA module using Gene Ontology (GO) and Kyoto Encyclopedia of Genes and Genomes (KEGG) pathways using the kegga() function from the limma package. Top GO and KEGG terms were extracted using topKEGG() with *p*-value cutoff of 0.001, as raw *p*-value were used without multiple-testing correction due to the hierarchical and non-independent nature of terms. While there are only 24 KEGG pathways for honeybees, there are 471 GO terms from in the three main hierarchical ontologies: Biological Process (BP), Molecular Function (MF) and Cellular Component (CC).

### Statistical Analysis

All data processing, normalization, statistical analysis and visualization were performed in R (version 4.5.0, 2025-04-11).

## Results

### Differential gene expression analysis

Based on one-way ANOVA with FDR < 0.05, total 9,345 genes (97.30%) showed a significant expression difference across low and high biting honeybees across different ages (Fig 1). The heatmap showed that distinct clustering of samples according to biting behavior and ages, indicating that both biting behavior and age influence gene expression patterns. Several clusters showed higher expression in high-biting honeybees across multiple age groups, while other clusters were highly expressed in low-biting honeybees.

**Figure 1.** Heatmap displaying 9345 genes with one-way ANOVA with FDR *p*-value < 0.05 across day 1 and day 8 worker honeybees with low and high biting behavior from Remove Unwanted Variation (RUV) model analysis.

### Pairwise Comparisons

Table 1 shows the number of significantly up-regulated and down-regulated genes for the pairwise comparisons and interaction test with traditional separate FDR correction for each comparison and *p*-value at 0.05. Comparison of LB and HB workers showed several differentially expressed genes. In workers collected on day 1 detected 166 up-regulated and 403 down-regulated genes, whereas workers collected on day 8 showed 82 and 46 up- and down-regulated genes, respectively. However, because multiple contrasts were examined, some of which had more expression changes than others, therefore, a “global” FDR across raw *p*-values with FDR < 0.05) for all contrasts together were performed (Supplementary Table 1). This ensures that a gene with the same raw *p*-value in two different comparisons would not end up with vastly different FDR *p*-values. Specifically, comparing LB and HB workers, 341 and 578 down- and up-regulated genes, respectively were detected on day 1, whereas 261 down-regulated and 198 up-regulated genes were identified on day 8. Heatmap of top 30 differentially expressed genes associated with biting behavior across 1-day and 8-day old honeybees with low and high biting behavior

**Table 1.**
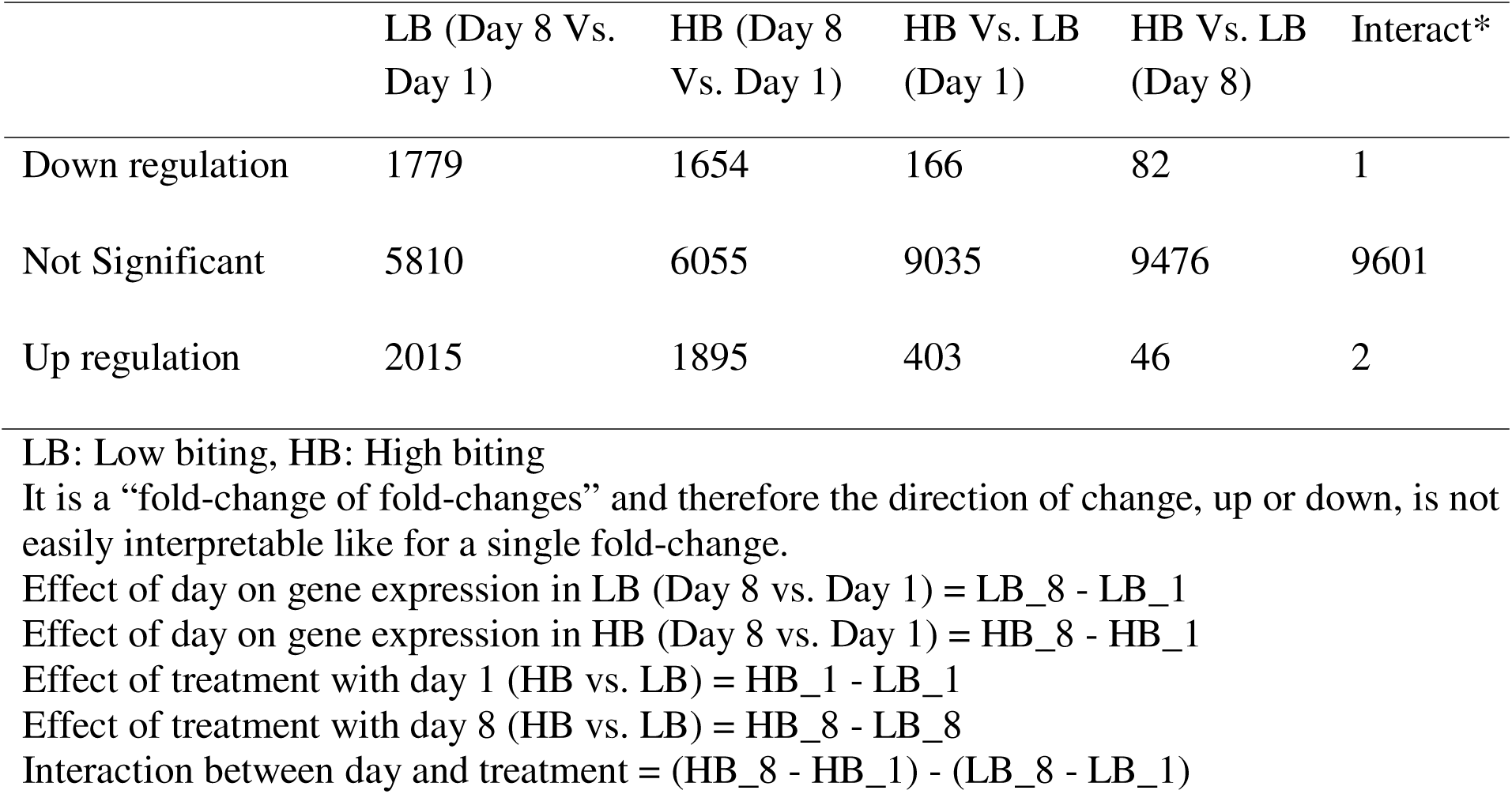
Differentially expressed genes (DEGs) in high and low biting honeybees collected on day 1 and day 8 (FDR < 0.05 by separate correction)

To identify high-confidence candidate genes associated with biting behavior, top 30 DEGs were filtered based on adjusted *p*-values, fold-change magnitude, and consistency cross contrasts statistical significance across day 1 and day 8 worker bees with low and high biting behavior (Fig 2). Several candidate genes identified in biting honeybees are associated with neural signaling, metabolic regulation and sensory responsiveness including *CG30101* (also known as *Vajk-4* gene), *yellow-y*, 409422, and potassium channel-related genes, suggesting potential involvement of neurobehavioral pathways underlying defensive biting behavior. The DEGs also included several uncharacterized LOC genes, viral transcripts or stress-associated responses, such as Deformed wing virus (DWV) or Varroa destructor virus (VDV).

**Figure 2.** Heatmap of top 30 differentially expressed genes associated with biting behavior across day 1 and day 8 worker honeybees with low and high biting behavior. Scaled fold-change values are shown, and hierarchical clustering reveals distinct expression modules linked to behavioral state and developmental regulation.

Differential expression patterns of high biting and low biting honeybees collected on different days including all up-regulated, down-regulated and not significant genes were also visualized using a mean-difference (MD) plot based on pairwise comparisons (Supplementary Figure 1 A, B, C, D). Despite the changes in total counts, the overall patterns of DGEs remained consistent, indicating that the main outputs are valid to the choice of multiple-testing correlation methods.

### WGCNA analysis

WGCNA was performed to identify modules of co-expressed genes associated with LB and HB behavior across different ages in honeybees (Fig 3 A, B, C, D and Supplementary Table 2). The WGCNA resulted in 10 modules ranging from 3305 to 107 genes (Supplementary Table 2). A total of 284 genes were assigned to module 0; however, eigengene values for this module were not analyzed because they do not represent coherent co-expression relationships (Supplementary Table 2). The brown module with 904 genes showed significantly higher expression in HB worker honeybees compared with LB worker honeybees on both day 1 (FC = 1.52, FDR = 0.0019) and day 8 (FC = 1.27, FDR = 0.056). Similarly, the yellow module that contains 563 genes also demonstrated strong behavioral associations, with higher expression in HB worker honeybees on both day 1 (FC = 1.47, FDR = 2.7 × 10^-4^) and day 8 (FC = 1.16, FDR = 0.073). In contrast, the green module including 550 genes consistently exhibited lower expression in HB worker honeybees relative to LB worker honeybees at both day 1 (FC = 2.3 ×10^-5^) and day 8 (FC= −1.47, FDR = 2.3 × 10^-5^). The turquoise module contains 3305 genes and displayed significantly upregulated on day 8 compared with day 1 in both LB (FC = 1.48, FDR = 1.2 × 10^-5^) and HB worker honeybees (FC = 1.49, FDR = 1.2 × 10^-5^).

**Figure 3.** Weighted Gene Coexpression Network Analysis (WGCNA) analysis across day 1 and day 8 worker honeybees with low and high biting behavior. Heat map of the respective module with normalized log_2_ expression for different genes across all samples and the bar plot of module eigengene expression (± SE) for day a vs day 8 for high and low biting behavior honeybees. The heat map and bar plot of (A) Brown module (B) Yellow module (C) Green module and (D) Turquoise module.

### Functional enrichment analysis

Functional enrichment analysis associated with biting behavior was conducted using GO and KEGG analyses. GO enrichment analysis showed over-representation of biological processes involved in neural signaling, sensory perception, muscle contraction, immune response, cellular respiration, metabolic processes, and structural organization (Fig 4 A, B, C, D, E). For example, the GO enrichment analysis of the genes in the brown module were significantly enriched in mitochondrial localization and metabolic processes. Similarly, cellular component analysis showed prominent involvement of mitochondrial and intracellular structures including mitochondrion, mitochondrial inner membrane, and organelle membrane. Biological processes primarily associated with metabolic processes such as carboxylic acid metabolic process, small molecule metabolic process, and primary metabolic process. Further, molecular function analysis revealed significant enrichment for catalytic activity and structural molecule activity, indicating their roles in metabolism and protein synthesis. The genes in the yellow module were significantly enriched in mitochondrial-related processes. Biological process included oxidative phosphorylation, electron transport chain, aerobic respiration, and ATP biosynthetic process. Cellular component analysis showed strong enrichment for mitochondrion, mitochondrial inner membrane, and mitochondrial protein-containing complex, indicating a central role of mitochondrial energy metabolism. Similarly, genes in the green module were significantly enriched in processes related to cell cycle regulation and genome maintenance. Biological processes include cell cycle, mitotic chromosome condensation and DNA damage response. The turquoise module revealed that its genes are significantly associated with biological processes such as cellular response to stress, DNA repair, and DNA damage response, indicating activation of stress-adaptive and genome maintenance mechanism. Cellular component enrichment included the endomembrane system, spliceosomal complex, and proteasome complex, while molecular functions were enriched in iron ion binding.

**Figure 4.** Gene Ontology (GO) enrichment analysis showing significantly enriched terms across biological process (BP), molecular function (MF), and cellular component (CC) among selected genes.

KEGG pathways analysis identified significant enrichment of various pathways including oxidative phosphorylation, cytoskeleton-related pathways, motor proteins, ribosome-associated pathways, citrate cycle (TCA cycle), and carbon metabolism which shows enhanced mitochondrial energy production associated with defensive responses (Fig 5 A, B, C). These pathways were more prominent on day 8 that associated with enhanced biting activity.

**Figure 5.** Kyoto Encyclopedia of Genes and Genomes (KEGG) pathway enrichment analysis illustrating significantly enriched biological pathways associated with the selected gene sets.

The brown module shows the ribosome pathway (44 DEGs/158 genes *p* = 2.67E^-07^) and carbon metabolism pathway (23 DEGs/87 genes, *p* = 0.00032) were among the top enriched pathways associated with the biting behavior in honeybee. The yellow module genes were primarily enriched in pathways related to energy metabolism and protein synthesis, including oxidative phosphorylation (55 DEGs/88 genes, *p* = 4.17 × 10^-40^), citrate cycle (TCA cycle) (21/32, *p* = 3.27 × 10^-16^), metabolic pathways (117/859, *p* = 1.83 × 10^-13^), carbon metabolism, ribosome, and motor proteins. . KEGG pathway enrichment analysis of turquoise module identified several pathways significantly associated with the differentially expressed genes. The proteasome pathway (29 DEGs/39 genes, *p* = 7.23E^-07^), biosynthesis of nucleotide sugars (18 DEGs/27 genes, *p* = 0.00044).

## Discussion

Worker honeybees undergo age-dependent shifts in task performance with newly emerged workers perform in-hive tasks, while older individuals gradually transition to defense and foraging. This age-dependent division of labor is closely linked with physiological and molecular changes that regulate behavioral maturation. Grooming behavior, including the physical biting and removal of mites, is an important component of honeybee defense against *Varroa* mite infestation (Li-Byarlay et al., 2025; Morin & Giovenazzo, 2023). Typically, defensive behavior in *A. mellifera* started after day 6 of emergence (Mondet et al., 2015; Russo et al., 2022). This study investigated the transcriptomic variation underlying high and low mite biting behavior in honeybees collected on day 1 and day 8 using differential gene expressions, co-expression network analysis and functional enrichment approaches. The findings of this study provide molecular insights into the complex behavioral and physiological mechanisms underlying mite-biting behavior in honeybees.

Age is a major determinant of physiological and behavioral state in honeybee workers, and previous transcriptomic studies have demonstrated profound age-dependent shifts in gene expression profiles associated with behavioral maturation and division of labor (Hamilton et al., 2019; Khamis et al., 2015; Whitfield et al., 2006). Age-related transcriptomic studies in honeybees have shown that genes involved in metabolic processes neural signaling, immune responses, sensory perception, energy metabolism and hormonal regulation change markedly as workers transition from in-hive tasks to aggressive behaviors or defensive roles (Chandrasekaran et al., 2015; Chen et al., 2025; Evans et al., 2006; Holt et al., 2013). Consistent with these findings, the present study revealed distinct sets of age-associated DEGs between high- and low-biting individuals, suggesting that biting behavior may be modulated by age-dependent molecular mechanisms. Differences in age-specific gene expression between HB and LB honeybees suggest that age interacts with behavior-specific regulatory pathways to shape biting behavior (Mu et al., 2025). The clustering top 30 DEGs included functional categories related to neural signaling, metabolic regulation and sensory responsiveness suggest that biting behavior is associated with coordinated regulation of metabolic and neural signaling pathways. For example, CG30101 (also known as *Vajk-4* gene) that is associated with cuticular protein involved in chitin-based cuticle development and denticle morphogenesis (Cinege et al., 2017). Furthermore, the presence of numerous uncharacterized genes among DEGs suggest that additional, currently unknown molecular pathways may contribute to defensive biting behavior. Further functional characterization of these unannotated genes may improve understanding of the genetic mechanisms associated with defensive and mite biting behavior in honeybees. These findings suggest that biting behavior in honeybees is associated with widespread transcriptomic changes rather than isolated gene effects, providing a foundation for further functional analyses.

Functional enrichment analysis through GO showed enrichment of several pathways associated with biting behavior in honeybees. Behavioral transitions in honeybees are regulated by the differential expression of multiple molecular pathways including juvenile hormone signaling, vitellogenin pathway, neural signaling, energy metabolism, and octopamine signaling (Khamis et al., 2015; Schilcher & Scheiner, 2023; Shpigler et al., 2017). Furthermore, previous studies also highlighted the role of transcription factors, neural signaling, sensory perception, metabolic processes and hormone-related pathways in regulating defensive or grooming behavior in honeybees (Alaux et al., 2009; Chandrasekaran et al., 2015; Hamiduzzaman et al., 2017; Holt et al., 2013). In the present study, functional enrichment analyses identified several biological processes that may be associated with mite biting in honeybees, including neural and sensory associated signaling, chitin binding, mitochondrial energy metabolism, stress response and neuromuscular function. Enrichment of neural-related processes such as signal transduction, axon guidance, neuron development and receptor-mediated signaling indicate that differences in biting behavior are associated with modulation of neural circuit activity and sensory processing (Alaux et al., 2009; Hunt, 2007). Similarly, enrichment of chitin-binding process may associated with robust mandible and indicate involvement of structural and extracellular matrix components in biting and defensive interactions (Klunk et al., 2024; Krings et al., 2024). Furthermore, strong enrichment of mitochondrial pathways suggests that biting behavior is associated with elevated energetic demands of metabolic process (Chandrasekaran et al., 2015; Li-Byarlay et al., 2014; Rittschof et al., 2018). In addition, the enrichment of muscle contraction and sarcomere organization pathways is associated with biting behavior in bees and other insects (de Meira & Gonçalves, 2021; Paul, 2001). The involvement of stress response including DNA repair, protein degradation, and proteasome further shows that biting behavior may associated with broader physiological stress adaptation (Even et al., 2012). These findings show that mite biting behavior is not driven by a single pathway but rather emerges from an integrated physiological process associated with neural excitability metabolic activation, cellular stress regulation and motor function (Li-Byarlay et al., 2014; Traniello et al., 2023).

KEGG exhibited the enrichment of metabolic and mitochondrial pathways in honeybees exhibiting high biting behavior indicate that biting behavior defensive is linked to energetic capacity and muscular function (de Meira & Gonçalves, 2021). Biting behavior requires rapid and sustained muscle contraction, which is supported by increased oxidative phosphorylation and central carbon metabolism (Li-Byarlay et al., 2014). Furthermore, enrichment of cytoskeletal and motor protein pathways suggests structural adaptation of muscle cells to facilitate aggressive responses. The stronger enrichment observed on day 8 compared to day 1 indicates that defensive behavior may be reinforced over time through transcriptional reprogramming, reflecting physiological maturation or behavioral plasticity in honeybees.

Overall, this integrative transcriptomic analysis reveals that biting behavior in honeybees is associated with age-dependent gene expression changes, behavior-specific regulatory pathways, and coordinated gene networks. These findings suggest that honeybee biting behavior is not governed by a single gene but rather by a coordinated network of metabolic, signaling, and structural genes that are modulated by both behavioral state and developmental stage. These findings contribute to a deeper understanding of the genetic and physiological basis of behavioral diversity in honeybee.

## Data availability statement

The raw reads of RNA-Seq related to this study are available in NCBI……

## Authorship contribution

H.L-B.: Conceptualization, Data curation, Funding acquisition, Methodology, Resources, Supervision, Validation, Writing—review & editing; M.C.F.: Conceptualization, Methodology, Investigation, Data curation; RP: Data curation, Data analysis, Writing—original draft. All authors reviewed the final manuscript.

## Supplementary materials

Supplementary material associated with this study are available in the online version of this paper.

## ACKNOWLEDGEMENTS

We acknowledge the Genomics Shared Resource (GSR) at OSUCCC-The James for their support with the library preparation and sequencing. GSR is funded in part by the National Cancer Institute P30 CA016058. This research was supported by USDA NIFA Evans-Allen funds NI241445XXXXG004, NI251445XXXXG001, USDA AFRI award 2020-67014-31557.

---

Supplementary Figure 1. Mean-difference plot with pairwise comparison between day 8 and day 1 honeybees collected from low and high biting colony. Each gene is plotted with its mean expression value on the X-axis and the log fold-change on the Y-axis. Genes with FDR p < 0.05 are colored either red (up-regulated) or blue (down-regulated). (A) high-biting colony (B) low-biting colony (C) Day 1 (D) Day 8

